# Elucidating the molecular interactions between uremic toxins and the Sudlow II binding site of human serum albumin

**DOI:** 10.1101/2020.02.19.940171

**Authors:** Josh Smith, Jim Pfaendtner

## Abstract

Protein bound uremic toxins (PBUTs) are known to bind strongly with the primary drug carrying sites of human serum albumin (HSA), Sudlow site I and Sudlow site II. A detailed energetic and structural description of PBUT interactions with these binding sites would provide useful insight into the design of materials that specifically displace and capture PBUTs. In this work, we used molecular dynamics (MD) simulations to study in atomistic detail 4 PBUTs bound in Sudlow site II. Specifically, we used the experimentally resolved X-ray structure of simulated indoxyl sulfate (IS) bound to Sudlow site II (PBD ID: 2BXH) to generate initial binding poses for p-cresyl sulfate (pCS), indole-3-acetic acid (IAA), and hippuric acid (HA). We calculated the interaction energy between toxin and protein in MD simulations and performed mean shift clustering on the collection of molecular structures from MD to identify the primary binding modes of each toxin. We find that all 4 toxins are primarily stabilized by electrostatic interactions between their anionic moiety and the hydrophilic residues in Sudlow site II. We observed transience in the strongest toxin-protein interaction, a charge-pairing with the positively charged R410 residue. We confirm the finding that the primary binding pose of IS in Sudlow site II is stabilized by a hydrogen bond with the carbonyl oxygen of L430, and find that this is also true for IAA. We provide insight into the chemical functional groups that might be incorporated to improve the specificity of synthetic materials for PBUT capture. This work represents a next step toward the *de novo* design of solutions to the problem of PBUT management in CKD patients.

**Significance Statement:** In spite of their implication in poor clinical outcomes, surprisingly little information is available about the structure and mechanisms that govern the binding of protein bound uremic toxins to their primary carrier human serum albumin. To date, only the structure of indoxyl sulfate has been determined by experiment. This paper describes a comprehensive characterization of four toxins that are known to bind Sudlow site II using molecular dynamics simulations. Based on the experimental structure of indoxyl sulfate bound to HSA, the binding mode within Sudlow site II of three additional PBUTs was determined. The structures, energetic and mechanistic analysis provide substantial new information for the nephrology community about these toxins as well as new protocols to aid future studies of PBUTs.

## 1. Introduction

The removal of protein bound uremic toxins (PBUTs) from the bloodstream of chronic kidney disease (CKD) patients is an unmet challenge. PBUTs are uremic retention solutes with a significant fraction of the toxin mass in the bloodstream existing in complex with proteins – primarily human serum albumin (HSA) – which renders them unsusceptible to clearance by traditional dialysis.^1,2^ Several of these toxins, including indoxyl sulfate (IS) and p-cresyl sulfate (pCS), have been correlated with poor clinical outcomes for CKD patients.^3–5^ Fully functional kidneys are surprisingly effective at managing PBUT levels in the bloodstream, but the molecular mechanisms for natural PBUT clearance have not been characterized.^6^ Without a detailed understanding of the protein unbinding and renal clearance process to guide design, strategies for artificial PBUT clearance in CKD have not produced comparable efficacy.

Several strategies have been suggested for managing PBUT concentrations in CKD patients, aimed at displacing toxins from their protein binding site (rendering them dialyzable) and more efficiently capturing free toxin from solution. A few of the most extensively studied (and clinically relevant) PBUTs are known to bind to the major drug binding sites of HSA, called Sudlow site I and Sudlow site II.^7^ Armed with this knowledge, Tao et al. recently demonstrated that introducing competitive binders for Sudlow site II (ibuprofen and tryptophan) to uremic plasma increased the unbound fraction of the uremic toxins IS, pCS, indole-3-acetic acid (IAA), and hippuric acid (HA).^8^ Others have demonstrated that diluting uremic plasma with hypertonic solutions, thereby increasing the ionic strength, facilitated PBUT unbinding and increased the clearance of IS and pCS during *in vitro* and *ex vivo* hemodialysis.^9,10^ The use of simple adsorbent chemistries targeting the hydrophobic and negatively charged moieties of many PBUTs has also shown promise in improving the efficiency of free PBUT capture.^11,12^ The efficacy of these strategies could be greatly increased given information about the dominant stabilizing interactions between PBUTs and the protein residues in their native binding sites.

Protein data bank structures of CMPF and IS bound to Sudlow site I and Sudlow site II, respectively, are the only molecular scale data currently available to describe the interactions between PBUTs and HSA at the molecular scale.^13^ While these experimental binding poses have provided valuable preliminary insights into the protein residues which stabilize CMPF and IS, they fail to capture the inherently dynamic nature of the protein-ligand complex. Also, insights derived from the experimental binding poses do not necessarily generalize to other PBUTs. Luckily, the 3-dimensional structures for CMPF and IS provide an entry point for more comprehensive characterization of PBUT-HSA interactions with molecular dynamics (MD) simulations. MD is a computational tool especially well-suited for investigating the inherently dynamic interactions of protein-ligand complexes with atomic resolution.

Sudlow site II is an attractive starting point for investigation with MD simulations because it has been established as the primary binding site of IS and pCS, the two PBUTs with the most extensive body of literature to support their deleterious physiological effects.^14^ The chemical similarity between IS and other Sudlow site II binders, including pCS, IAA, and HA, can also be exploited to develop initial binding poses for MD simulations of other toxins. **Figure 1(a)** shows the chemical structures of the 4 toxins IS, pCS, IAA, and HA. **Figure 1(b)** shows a 2D PoseView representation of the only experimentally resolved crystal structure of a PBUT (IS) bound to Sudlow site II (PDB ID: 2BXH).^13^

**Figure 1.**
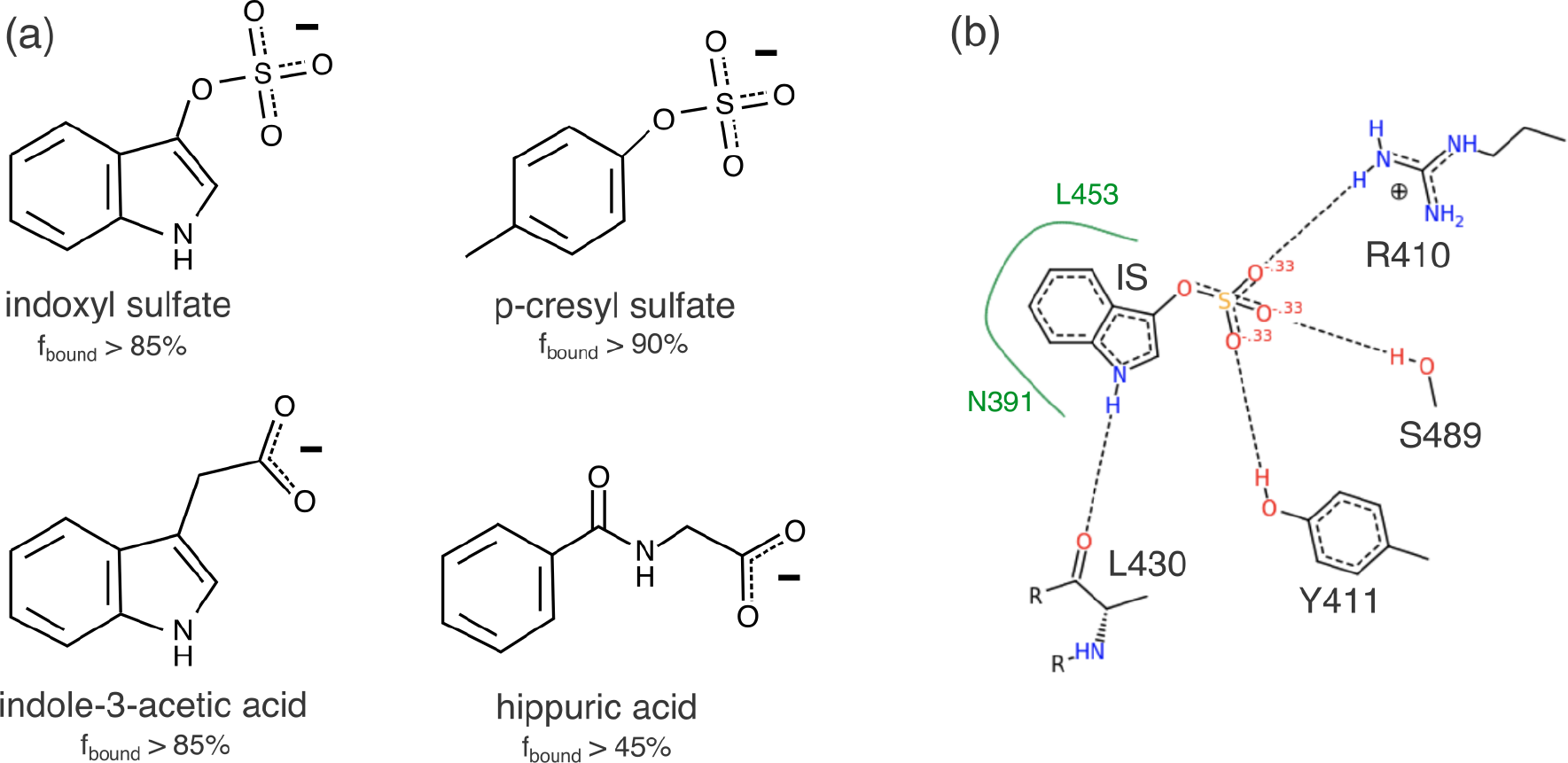
Two-dimensional representations of PBUTs known to bind to the Sudlow site II of HSA with high affinity. (a) Chemical structures of uremic retention solutes known to bind to the Sudlow site II of human serum albumin. The fraction of toxin that exists bound to HSA (f_bound_) in the bloodstream is included for each toxin, according to reference 14. (b) PoseView representation of the experimentally resolved binding structure for indoxyl sulfate in Sudlow site II (PDB: 2BXH). Putative hydrogen bonds are represented by dashed gray lines, and hydrophobic interactions by solid green lines.

In this article, we describe an extensive molecular dynamics (MD) simulation study of PBUTs in the Sudlow site II of HSA. We simulated the IS-HSA complex in water, starting from the experimental binding pose. We performed identical simulations with pCS, IAA, and HA in place of IS, starting with initial binding poses derived from the IS-HSA binding pose. The energetics of the protein-toxin interactions were used to identify the key toxin-stabilizing residues in Sudlow site II. Unique binding modes for IS, pCS, IAA, and HA were identified with unsupervised learning.

Our analysis provides insight into the general features of PBUT binding in Sudlow site II, as well as insights into toxin-specific interactions. All 4 toxin residues were found to be primarily stabilized by electrostatic interactions with hydrophilic protein residues. The primary binding modes for IS and IAA were stabilized by a hydrogen bond with the carbonyl oxygen of L430 deep in the hydrophobic core of the binding pocket, while pCS and HA were stabilized by hydrogen bonding interactions near the mouth of the binding pocket. Excellent agreement was observed between the experimental binding pose of the IS-HSA complex and the primary binding mode identified with MD, providing external validation for our results. This work represents a new contribution to the molecular level understanding of PBUT-HSA interactions and establishes a baseline for more advanced molecular modeling studies in the future.

## 2. Methods

### 2.1 Preparing initial structures for HSA-toxin complexes

Of the uremic toxins known to bind Sudlow site II with high affinity, an experimentally resolved structure of the toxin in complex with HSA has only been reported for indoxyl sulfate (RCSB PDB ID: 2BXH).^13^ The 2BXH PDB structure contains an HSA dimer, each associated with 3 IS molecules associated and crystallographic waters. We took the coordinates of the “chain A” HSA molecule (atoms 1-4263) and the IS molecule bound to its Sudlow site II (atoms 8531-8544). The other 2 IS molecules associated with chain A, overlapping reconstructions of alternative binding poses of IS to Sudlow site I, were ignored based on experimental evidence that suggests there is negligible IS binding to Sudlow site I under physiologically relevant conditions.^15^ The tleap program from Amber tools was used to add hydrogens and missing atoms to the HSA molecule.^16^ We built an IS molecule (complete with hydrogens) in GaussView and superimposed this upon the IS heavy atoms from the PDB using the “RMSD Calculator” extension in VMD.^17,18^ Gromacs tools were used to construct a cubic box (with sides 10.8 nm in length) around the IS-HSA complex and to add TIP3P water and sodium counter ions.^19,20^ The Amber 14 force field parameters were used for the HSA protein.^21^ We calculated the partial charges for IS atoms using the restrained electrostatic potential (RESP) method and assigned bonded parameters according to the Generalized Amber Forcefield (GAFF).^22,23^ Quantum mechanical calculations for RESP were performed with Gaussian 09 using density functional theory with the B3LYP/6-31G* level of theory.^18^

Motivated by the chemical similarity between IS and the other uremic that bind to Sudlow site II, we used the IS-HSA PDB structure as a template for putative binding poses of p-cresyl sulfate (pCS), indole-3-acetic-acid (IAA), and hippurate. We built each toxin molecule in GaussView and superimposed the structure onto the IS-HSA structure in VMD so that the anionic and hydrophobic moieties of the new toxin aligned with the sulfate and indole groups of the IS molecule, respectively. The initial superimposed structures are pictured in **Figure S1**. Solvent and counter ions were added to each of these structures following the method outlined above for IS. An identical protocol was also used for assigning forcefield parameters for each toxin.

### 2.2 Molecular dynamics simulations of the toxin-HSA complexes

All molecular dynamics (MD) simulations were performed using Gromacs 2016.3.^19^ A preliminary 3-step equilibration protocol was used to relax the initial solvated structure of each protein-toxin complex, which included energy minimization, annealing, and a short simulation in the NPT ensemble. Energy minimization was performed on the initial solvated structures following the steepest descent algorithm for 10000 steps. All other simulations were propagated using an integration timestep of 2 fs. A 250 ps annealing simulation was performed to gently relax the system from the energy minimized conformation to a realistic configuration at the target temperature of 298 K. This was achieved by ramping the coupling temperature for the Bussi-Donadio-Parrinello thermostat smoothly from 5 to 298 K over the first 240 ps of the simulation.^24^ The output configuration and velocities from the annealing simulation were used to initiate a 500 ps NPT simulation with the Berendsen barostat for rapid coupling to the target pressure of 1 bar.^25^

The strong binding affinity between HSA and the 4 PBUTs suggests that unbinding events are not likely to occur on the timescale currently accessible with unbiased molecular dynamics simulations. We added an additional equilibrium simulation to ensure that each protein-toxin complex (especially the pCS, IAA, and hippurate complexes) reached a metastable binding pose that would not lead to rapid unbinding in production simulations. We performed a 5 ns NPT simulation with the Bussi-Donadio-Parrinello thermostat and Parrinello-Rahman barostat.^24,26^ The system temperature was coupled to 298 K with a frequency of 10 ps^−1^, and the pressure was coupled to 1.0 bar with a frequency of 1 ps^−1^. The potential energy of the system was extended with an upper wall potential, implemented with Plumed 2.4,^27^ according to the following

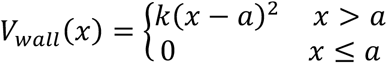

where x is equal to the root mean square deviation (RMSD) of carbon atoms of the toxin and neighboring binding pocket residues, a is the RMSD value at which the restraining potential is turned on (in this case 0.1 nm), and k is the restraining force constant (in this case 150 kJ/mol-nm^2^). This restraint was intended to prevent rapid unbinding as might occur with a poor initial guess at the binding pose, which could result in steric clashes or atomic overlap.

Three 250 ns NPT simulations were performed for each protein-toxin complex (12 production simulations in total). Each production simulation was initiated with the output configuration from the restrained binding-pocket simulation and different randomly generated velocities. The Bussi-Donadio-Parrinello thermostat and Parrinello-Rahman barostat were used to maintain a temperature and pressure of 298 K and 1 bar, respectively.^24,26^ The first 50 ns of the simulation were allowed for any further relaxation of the protein structure, and the final 200 ns of each simulation were used for all following analysis.

### 2.3 Energy and structural descriptor calculations

Protein-toxin interaction energy was calculated using the GROMACS tool ‘gmx energy’. The initial list of 37 residues to be screened for interactions with each toxin was generated by taking all protein residues with at least one atom within 6 Å of any atom of indoxyl sulfate in the experimentally resolved PDB structure (PDB: 2BXH). The number of atomic and hydrophilic contacts between each of these residues and each toxin were calculated with Plumed 2.4 using the famous Bonomi equation, a switching function defined as follows

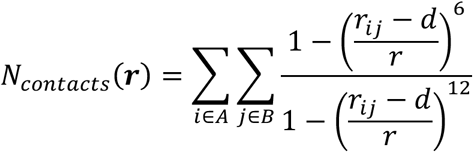

where *A* and *B* represent the sets of heavy atoms (nonhydrogen) atoms of the toxin and a given protein residue, *r*_*ij*_ is the intermolecular distance between atoms *i* and *j*, and *d* and *r* are switching function parameters that control the diameter and radius of a coordination shell around each atom. Atomic contacts were calculated every 10 fs (5 MD steps) of each production simulation, setting a 4 Å coordination sphere around each atom (r = 0.4 nm, d = 0). Hydrophilic contacts were calculated with a coordination annulus around each atom, 2.7 Å in diameter and 0.3 Å thick (r = 0.03 nm, d = 0.27 nm). This definition of hydrophilic contacts is consistent with the donor-acceptor distance expected for strong hydrogen bonds.

For the 9 residues selected for further analysis (selected based on atomic and hydrophilic contacts), the Lennard-Jones and Coulomb components of interaction energy with the toxin were calculated every 2 ps of each production simulation, again using ‘gmx energy’. The center of mass distance between each of the 9 key residues and the toxin (and the minimum distance between any R410 heavy atom and the toxin heavy atoms) were calculated every 2 ps of each simulation using Plumed 2.4. Subsequent analysis and visualization of these energetic and structural descriptors were performed in the Python programming language. Data files and analysis scripts have been made publicly available at www.github.com/UWPRG/pbut_analysis.

### 2.4 Dimensionality reduction and clustering with principal component analysis (PCA) and mean shift

With the large number of MD frames (300,000 for each complex) and the relatively high dimensionality of the structural descriptors for each frame (9 center of mass distances), an automated solution for pattern recognition was required to identify different binding states. We used the mean shift algorithm for clustering. Mean shift is a mode-seeking algorithm that essentially identifies peaks (modes) in a probability density function and clusters points that climb to the same peak following gradient ascent.^28^ This results in a more physically meaningful clustering than other commonly used clustering algorithms such as k-means (see **Figure S2** for further explanation). The natural fit of mean shift for processing data from biophysical simulations has been previously noted.^29,30^

The results of mean shift clustering are dependent on an initial estimate of the underlying probability density. Nonparametric techniques for density estimation suffer from the curse of dimensionality and perform better on dense, low dimensional data. A principal component analysis (PCA) was performed to project the 9 toxin-residue center of mass distances for each frame of the MD simulations onto a 2 dimensional structure space. PCA is a dimensionality reduction technique that is commonly used to project the inherently high-dimensional data from molecular dynamics simulations into an informative low-dimensional space. The coefficients for each principal component are provided in **Table S1**. We found that considering 3 or more principal components did not qualitatively change the clustering results. Mean-shift clustering was performed on the PCA-transformed data to identify unique metastable states in this dimensionally-reduced structure space for each protein-toxin complex. The average silhouette score – a quantitative measure to assess the quality of clustering – was used to select the optimal bandwidth for density estimation in the 2D principal component space (**Table S2**).^31^

## 3. Results and Discussion

We performed MD simulations of each PBUT-HSA complex in aqueous solution to investigate the key stabilizing interactions and structural features of the bound state. For each toxin we performed 3 classical MD simulations, each 250 ns long. The first 50 ns of each simulation was used to allow further relaxation of the toxin-protein complex. Frames were saved for analysis every 2 ps (1000 MD steps) over the final 200 ns of each replica. Unless otherwise stated, the average values for energetic and structural observables reported in **Section 3** were calculated from the aggregated frames of all 3 simulations and the standard deviations were calculated from the mean values for each replica (standard deviation over 3 means).

### 3.1 Uremic toxins are primarily stabilized by electrostatic interactions in Sudlow site II

Each of the PBUTs considered in this work is comprised of a bulky hydrophobic moiety and a hydrophilic, anionic moiety. As a first step in characterizing the nature of PBUT binding to Sudlow site II of HSA, we investigated the relative contributions of hydrophobic and electrostatic interactions to the overall protein-toxin interaction energy in the bound state. We calculated the interaction energy between the toxin and the protein in each frame of the simulation and tracked the contributions due to the Lennard-Jones, E_LJ_, and Coulomb, E_C_, potentials over time. The average E_LJ_ and E_C_, as well as the total interaction energy, for each complex are tabulated in **Table 1**.

**Table 1.** Average interaction energy between HSA and uremic toxins bound to Sudlow site II.

| | $E_C (\pm \text{sem})$<br>[kJ/mol] | $E_{LJ} (\pm \text{sem})$<br>[kJ/mol] | $E_{\text{total}} (\pm \text{sem})$<br>[kJ/mol] |
| --- | --- | --- | --- |
| Indoxyl sulfate (IS) | - 114 (6.12) | - 103 (1.68) | - 217 (7.38) |
| p-cresyl sulfate (PCS) | - 136 (8.92) | - 87.2 (0.138) | - 223 (9.08) |
| Indole-3-acetic-acid (IAA) | - 154 (8.37) | - 90.0 (0.403) | - 244 (8.21) |
| Hippurate (HA) | - 162 (27.3) | - 84.9 (3.75) | - 247 (24.5) |

For every toxin, the average magnitude of E_C_ is greater than that of E_LJ_. Not surprisingly, IS (the bulkiest of the 4 toxins) has the highest average Lennard-Jones interaction energy, followed by IAA. The bulky indole group (shared also by tryptophan, an endogenous HSA ligand and metabolic precursor to IS and IAA) apparently fits more tightly in the hydrophobic pocket of Sudlow site II than the smaller phenyl groups of pCS and HA.

Although IS and pCS have the same anionic moiety (sulfate) and IS has an additional hydrogen bond donor in its indole ring, pCS was found to have a lower average E_C_ than IS. We also found that, on average, pCS actually forms more hydrogen bonds with HSA residues in Sudlow site II than does IS (**Table S3**). This could be in part because the electron withdrawing N of the IS indole ring decreases the polarity of the sulfate linker oxygen atom of IS relative to the linker oxygen of pCS (see. mol2 files on GitHub). The carboxylate-containing toxins, IAA and HA, have stronger electrostatic interactions with the protein binding site than the sulfate-containing toxins, IS and pCS. This is likely due to the increased charge density of the carboxylate anion of IAA and HA compared to the sulfate anion of IS and pCS (see **Figure S3**). The geometry of the carboxylate anion also facilitates a very low-energy bidentate hydrogen bonding conformation with the guanidinium cation of the R410 residue. A representative structure for the HA-R410 double hydrogen bond conformation is provided in **Figure S4**.

The observation that average E_C_ is greater than average E_LJ_ in each case suggests that electrostatic interactions play an important role in stabilizing the protein-toxin complex. This is in good agreement with an experimental study that demonstrated the binding affinity between IS and HSA in various NaCl solutions decreased with increased ionic strength.^15^ Hypertonic predilution has been demonstrated to increase HD removal of only IS and pCS (as well as the uremic toxin phenyl acetic acid, which does not bind Sudlow site I or site II). The observation that all 4 PBUTs simulated in the present work are primarily stabilized by electrostatic interactions suggests that increasing ionic strength may be a general strategy for favoring PBUT dissociation from Sudlow site II.

The shape of the probability density function (pdf) for each component of the protein-toxin interaction energy provides additional information beyond the mean. **Figure 2** shows the probability distribution of E_LJ_ and E_C_ from all 3 simulations for each PBUT complex. An additional explanatory figure for **Figure 2**, showing the interaction energy vs. time for each MD simulation is provided in **Figure S5**.

**Figure 2.**
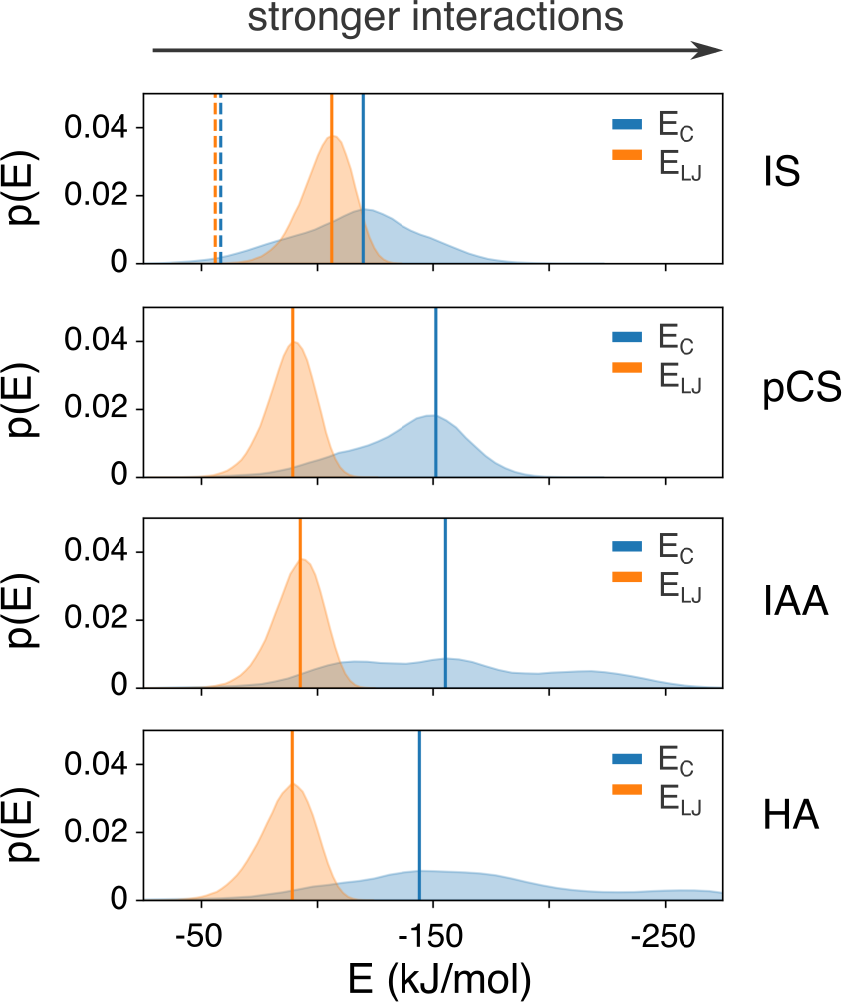
Probability distribution functions for interaction energies between toxins and protein residues calculated from MD simulations. The blue and orange shaded regions represent the pdf for E_C_ and E_LJ_, respectively, based on a histogram of every frame of the production MD simulations for each complex (3 × 100,000 observations). The peak of each pdf is marked by a solid, vertical line. The dashed, vertical lines in the top panel represent the E_C_ and E_LJ_ energies calculated from the experimental IS-HSA structure (PDB: 2BXH).

One would expect the pdf to have a single well-defined peak if fluctuations around a single low energy binding pose could account for the structural ensemble sampled with MD. Indeed, the pdf for the Lennard-Jones energy could be accounted for by a single peak. However, the broad irregular shape of the pdf for Coulomb energy for each complex suggests that each PBUT visits a number of metastable binding poses during the MD simulations (**Figure 2**). In **Sections 3.2-3.5**, we identify the key protein residues for stabilizing PBUTs in Sudlow site II and use structural descriptors derived from these residues to describe the metastable binding modes for each toxin.

### 3.2 Protein-toxin interactions are dominated by the same residues for all PBUTs tested

For further insight into the roles of specific Sudlow site II residues in stabilizing each toxin, and to identify possible structural distinctions between unique binding poses, we analyzed the interactions between toxins and binding pocket residues in more detail. We started with a list of 37 residues close to IS in the experimental xray structure, defining close residues as those with at least one atom (including hydrogens) within 6 Å of any atom of IS. To screen for residues of interest, we calculated the number of atomic and hydrophilic contacts between the toxin and each of these 37 residues in each frame of the MD simulations. The average number of atomic and hydrophilic contacts for each protein residue and each toxin are included in **Figure S6**.

Taking as the pertinent residues the set that includes all residues ranking in the top 5 residues in terms of either atomic or hydrophilic contacts for each toxin, we developed a list of 9 key residues for further analysis. A 3D structure highlighting the original 37 residues and filtered list of 9 residues is provided in the supporting information (**Figure S7**). This list includes all residues identified as hydrogen bond partners with IS in the experimental crystal structure (R410, Y411, L430, S489).^13^ The other key residues identified in this filtering process (L387, N391, K414, V433, L453) may help to distinguish between metastable binding modes for IS, or be involved in stabilizing the primary binding modes of the other toxins. **Figure 3** shows the average Coulomb and Lennard-Jones energy contributions of each key residue.

**Figure 3.**
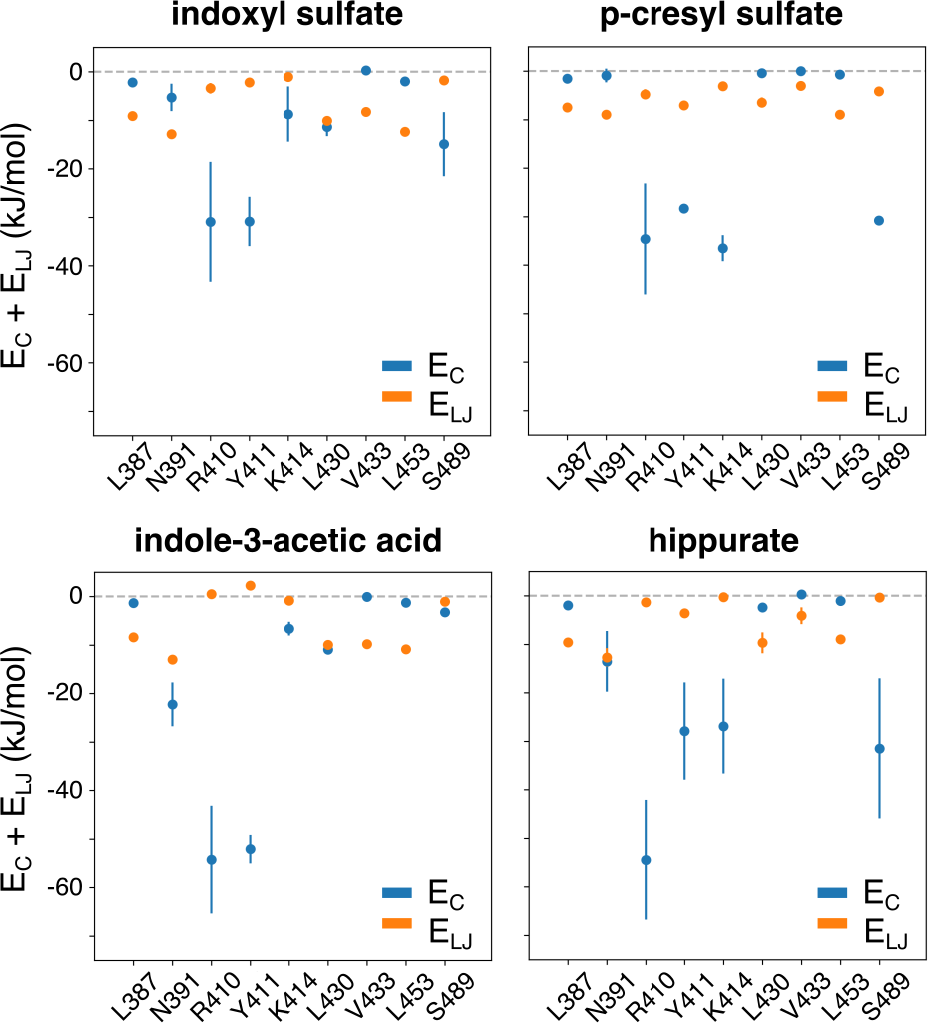
The interaction energies contributed by each of the 9 key binding pocket residues to the (left) IS-HSA complex and (right) pCS-HSA complex. The average Coulomb and Lennard-Jones components of the residue-toxin interaction energy (for all 3 simulations) are represented by blue and orange circles, respectively. Error bars represent +/− 1 standard deviation from the mean.

The two residues that interact most strongly with IS in the MD simulations are R410 and Y411. This observation is in good agreement with an experimental study that showed R410A and Y411A mutations together decreased the binding capacity of HSA for IS.^32^ R410 and Y411 are also the primary energetic contributors to the IAA complex. pCS and HA interact strongly with R410 and Y411, but also with K410 and S489, two other potential hydrogen bonding partners. These observations are particularly interesting because it might have been expected that the anion of the toxin would dictate its interactions with hydrophilic residues in the binding pocket.

Both IS and IAA participate in electrostatic interactions with the hydrophobic L430 residue. This interaction can be explained by a hydrogen bond between the indole NH group of each toxin and the carbonyl (backbone) oxygen of L430, pictured in the PoseView representation of the IS-HSA complex in **Figure 1(b)**. This interaction has previously been described as a polar feature inside the hydrophobic core of the binding pocket.^13^ Also of note, N391 contributes to the interaction energy with IS and pCS primarily through E_LJ_, but contributes also to the electrostatic interactions with IAA and HA.

### 3.3 Transient salt bridge with R410 is a primary contributor to low energy poses

The energetic contributions of R410 also have the largest standard deviation of the 9 residues. Large changes in the interaction energy between the toxin and one or a few residues may signify transitions between metastable states. To monitor the relationship between binding pocket conformation and interaction energy, we calculated the center of mass distance to each of the 9 key residues identified in **Section 3.2** and calculated their correlation with overall protein-toxin interaction energy. **Table 2** shows the Pearson correlation coefficient between the overall toxin-protein interaction energy and the center of mass distance between IS and each of the 9 residues.

**Table 2.** Correlation between proposed order parameters and protein-toxin interaction energy.

| indoxyl sulfate |  | p-cresyl sulfate |  | IAA |  | hippurate |  |
| --- | --- | --- | --- | --- | --- | --- | --- |
| Residue (ranked) | r | Residue (ranked) | r | Residue (ranked) | r | Residue (ranked) | r |
| R410 | 0.57 | R410 | 0.60 | R410 | 0.41 | R410 | 0.63 |
| Y411 | 0.44 | Y411 | 0.52 | Y411 | 0.30 | Y411 | 0.53 |
| S489 | 0.43 | S489 | 0.46 | L387 | 0.19 | K414 | 0.45 |
| K414 | 0.34 | K414 | 0.45 | N391 | 0.10 | S489 | 0.41 |
| L387 | 0.05 | L387 | - 0.13 | K414 | 0.07 | L453 | 0.00 |
| L453 | - 0.17 | N391 | - 0.26 | V433 | 0.01 | L387 | - 0.28 |
| L430 | - 0.19 | L453 | - 0.28 | L453 | 0.00 | V433 | - 0.34 |
| N391 | - 0.29 | L430 | - 0.42 | S489 | - 0.07 | L430 | - 0.39 |
| V433 | - 0.30 | V433 | - 0.44 | L430 | - 0.15 | N391 | - 0.56 |

Interestingly, there is a strong correlation between the protein-toxin interaction energy and the center of mass distance between IS and several of the residues (**Table 2**). There is a positive correlation between interaction energy and center of mass distance to each of the hydrophilic residues at the mouth of Sudlow site II, and a relatively strong negative correlation between interaction energy and the distance to hydrophobic residues buried in the binding pocket. This suggests an enthalpy gain for the toxins upon shifting away from the hydrophobic core and toward the opening of the binding pocket.

For each toxin, the R410-toxin distance has the strongest correlation with the interaction energy of the complex. Closer inspection of R410-toxin interactions showed that this important stabilizing residue frequently breaks contact with the toxin in favor of a fully solvated state. **Figure 4** shows that the breaking and forming of a salt bridge between IS and R410 is an important process in transitioning between high and low energy states.

**Figure 4.**
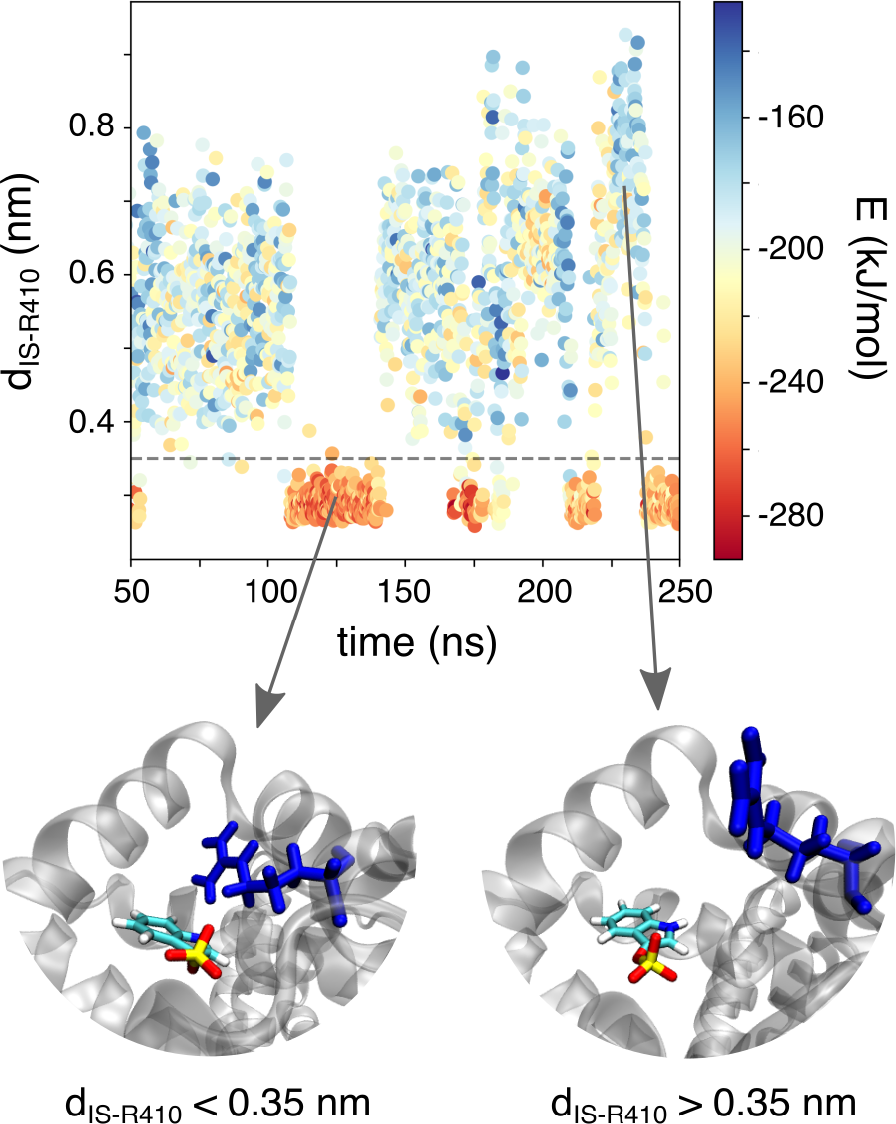
The minimum distance between IS and R410 heavy atoms vs. time from IS simulation 3, and its relation to the overall protein-toxin interaction energy. Each point is colored based on the overall protein-toxin interaction energy, ranging from dark red (lower energy) to dark blue (higher energy). The dashed horizontal line at 0.35 nm represents a putative cutoff for IS-R410 hydrogen bonding. The images below the plot show representative conformations for (left) low energy, low minimum distance and (right) high energy, high minimum distance.

**Figure 4** shows clearly that the IS-HSA complex is stabilized by salt bridging with the charge-neutralizing arginine in Sudlow site II. The weakened protein-toxin interactions associated with the breaking of this salt bridge, as well as the alleviation of potential steric hindrance to unbinding – observable in the representative conformations in **Figure 4** – signal the potential importance of this molecular movement in the unbinding process. The breaking/forming of this charge pair was observed multiple times for all toxins (**Figure S8**). Supporting the previous hypothesis that the charge-dense carboxylate groups provide especially strong interactions with R410, we find that the mean distance of R410 hydrogen bonds with IAA and HA is lower than those with IS and pCS (**Figure S8**).

Shifting the conformational equilibrium for R410 toward the solvated state could be a potential target for strategies to encourage toxin unbinding. The equilibrium could be shifted by adding cosolutes that lower the free energy of solvation for the R410 guanidinium side chain. R410 solvation could potentially be involved in the mechanism for IS release in hypertonic solutions. One could also target allosteric regulation of the Sudlow site II that specifically favors HSA conformations with R410 in the solvated state. Further investigation, both experimental and computational, into the effect on R410 position of ions in solution and remote binding to HSA could provide more actionable suggestions for toxin displacement strategies.

### 3.4 The toxin hydrophobic moiety determines the primary binding mode in Sudlow site II

The relatively fast breaking and forming of the R410 salt bridge can happen in isolation without significant movement of the bound toxin relative to other residues in the binding site. We performed a principal component analysis (PCA) on the 9 center of mass distances to project onto a low-dimensional conformation space where identifying unique states with unsupervised learning was more tractable. We used the mean shift algorithm to assign each frame of the MD simulation to a binding mode, based on the first two principal component values for the frame. A representative structure for each mean shift mode was taken as the MD frame with principal components nearest (based on 2D Euclidian distance) the center of that mode. **Figure 5(a-b)** shows the 2D probability density function along the first two principal components for the IS-HSA complex and the results of clustering points on this pdf with the mean shift algorithm. **Figure 5(c-f)** contains a summary of the structural features and representative structures of the IS binding modes identified with mean shift. Similar figures for the other 3 toxins are provides in **Figures S9-S11**.

**Figure 5.**
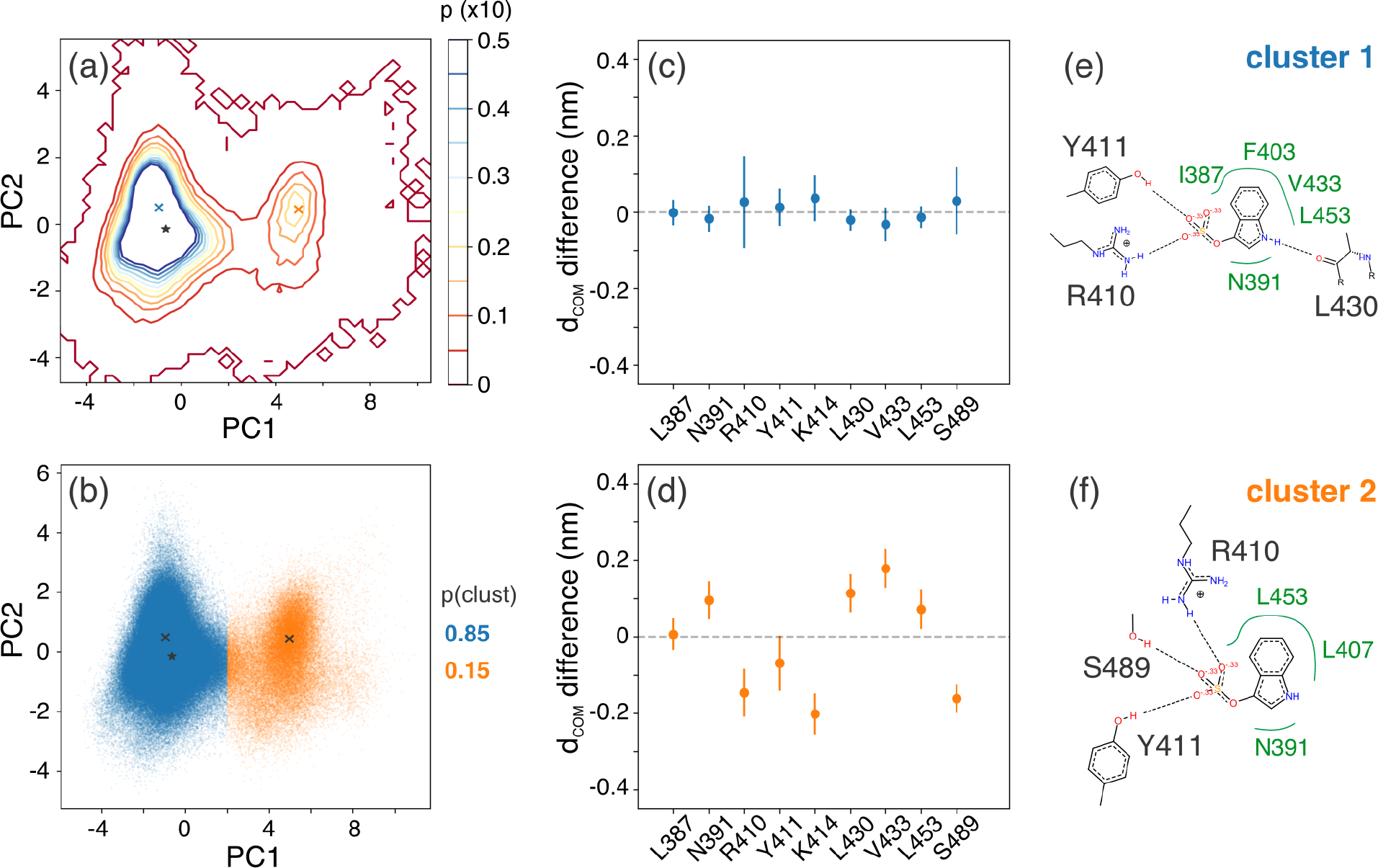
PCA and clustering results for the indoxyl-sulfate-HSA complex. **(a)** Contour plot of the 2D pdf for the IS-HSA complex, created from a 2D histogram of the PC values for each frame of the MD simulations, with a probability step of 0.005 between contour lines. Blue and orange ‘x’ symbols mark the center of each mode identified with mean-shift. The star symbol represents the experimental structure for IS-HSA (PBD: 2BXH). **(b)** Scatter plot of the first 2 PC values for each MD frame, colored by the cluster assigned by mean-shift. **(c-d)** The difference between the average IS-residue center of mass distance for all points in each cluster and the overall average IS-residue center of mass distance (all points in all clusters). **(e-f)** A PoseView representation of the central structure of each cluster (the MD frame with PC values nearest the ‘x’ symbols in **(a-b)**.

The first two principal components of the IS-HSA structural descriptors accounted for 57% and 15%, respectively, of the variance in the data set (PC coefficients available in **Table S1**). We identified 2 binding modes for IS in Sudlow site II, which accounted for fractions of 0.85 and 0.15, respectively, of the conformations sampled with MD. Each binding mode from MD has partial overlap with the experimental binding pose (**Figure 1b**). The primary binding mode for IS from MD and the experimental binding pose both have hydrogen bonding interactions with L430, R410, and Y411 and hydrophobic interactions with N391 and L453. Relative to the experimental structure, the primary MD structure is shifted deeper into the hydrophobic core of the binding pocket, interacting with hydrophobic residues I387, F403, and V433, and preventing a hydrogen bond with S489. In the secondary binding mode of IS, losing the L430 hydrogen bond has apparently allowed IS to shift nearer to the hydrophilic mouth of the binding pocket. In the secondary binding mode, IS regains the hydrogen bond with S489, but loses hydrophobic interactions with I387, F403, and V433. Further segmentation of these clusters, using kmeans clustering (k > 2), captures the movement of individual protein residues rather than significant changes in IS position (**Figure S12**).

The results of PCA and clustering for the other three PBUT complexes shed interesting light on the residue-wise energetic analysis described in **Section 3.2**. **Figure 6** shows a PoseView representation of the central structure of the most populated cluster for each complex.

**Figure 6.**
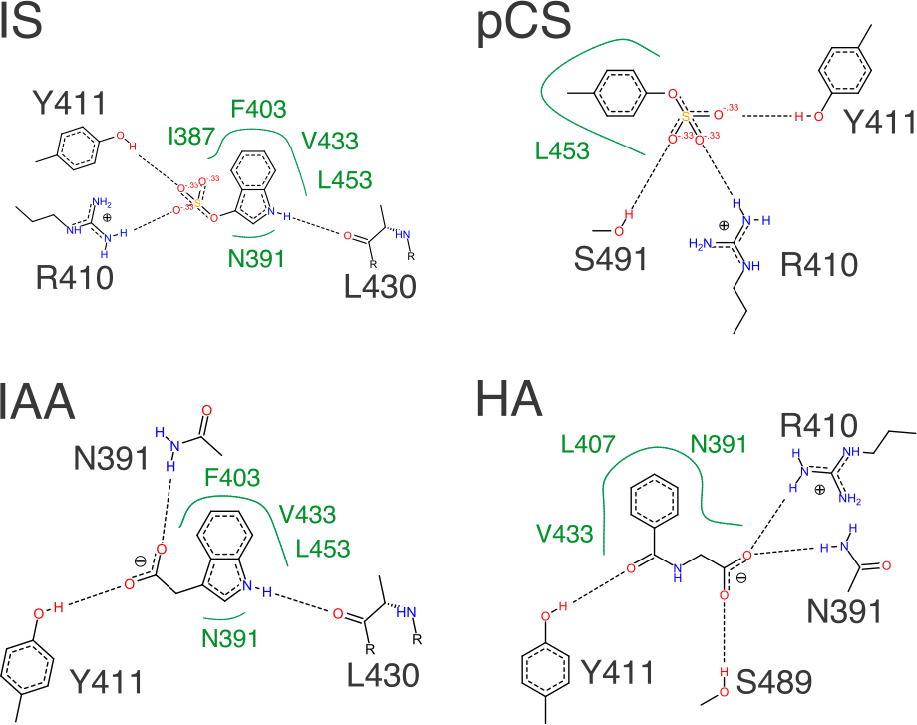
PoseView representation of the most populated binding mode for each PBUT-HSA complex. Putative hydrogen bonds are represented by black dashed lines, accompanied by the name and sidechain structure for the associated protein residues. Hydrophobic interactions are represented by solid green curves, labeled with the names of the associated protein residues. Polar atoms O, N, and S, are colored red, blue, and yellow, respectively.

The primary binding mode for IAA is also stabilized by a hydrogen bond with the L430 carbonyl oxygen, accounting for the electrostatic interactions with L430 identified in **Section 3.2**. The bulky indole groups of IS and IAA are apparently pinned deep in the hydrophobic core by this hydrogen bond and extensive hydrophobic interactions with other residues. Notably, only one binding mode was identified for IAA. The representative binding pose extracted from the sole IAA cluster captures a pose without hydrogen bonds donated from R410. As discussed in **Section 3.2**, the R410 position is observed to fluctuate in and out of hydrogen bonding distance with one, or both, of the carboxylate oxygens of IAA. The stability of the IAA binding mode suggests that hydrogen bonds with L430, Y411, and N391, can effectively hold IAA in place when the transient salt bridge with R410 is broken.

pCS and HA assume primary binding modes similar to the secondary binding mode of IS. Lacking an appropriately positioned hydrogen bond donor to exploit the L430 “polar” moiety, pCS and HA are not pinned in the hydrophobic pocket like IS and IAA. pCS and HA are allowed to shift closer to the mouth of the binding pocket where they interact with K414 and S489, as shown in **Section 3.2** (**Figure 3**). The carboxylate-containing toxins, IAA and HA, both accept hydrogen bonds from the NH_2_ group of N391 in their primary binding pose. This interaction is also observed in an alternate binding mode of pCS (**Figure S9**).

## Conclusions

The preference of each toxin for interactions with different chemical functional groups in Sudlow site II could be useful in designing toxin capture materials with high affinity and specificity for each toxin. Generally, a positively charged moiety, such as the guanidinium moiety of R410, and hydrogen bond donors, such as the hydroxyl of Y411 and S489, could be incorporated into synthetic constructs to nonspecifically stabilize all 4 toxins. Incorporating hydrophobic constituents into a synthetic construct for PBUTs may also be generally effective. Embedding a hydrogen bond acceptor, such as the L430 carbonyl, in the hydrophobic region may increase specificity for IS and IAA. The observation that a polar moiety embedded in a hydrophobic pocket stabilizes the primary IS binding mode may serve as a preliminary mechanistic explanation for the recent finding that cyclodextrin can bind IS from solution.^33^

Future work should target a direct connection between molecular modeling and the macroscopic observables that impact CKD patient health. Advanced simulation techniques for quantifying ligand binding affinity and unbinding kinetics represent promising avenues to make this connection.^34,35^ The current work provides the key ingredients necessary to use these advanced techniques, including representative structures for metastable binding poses and insight into discriminative structural features. Hopefully this contribution marks the beginning of a more precise and informed materials design process that will ultimately lead to better outcomes for CKD patients.

## Supporting Information

The SI contains additional figures and tables that have been described in the text above. To facilitate scientific reproducibility data files and analysis scripts have been made publicly available at www.github.com/UWPRG/pbut_analysis.

## Acknowledgements

The authors gratefully acknowledge a grant from the Northwest Kidney Centers to the Center for Dialysis Innovation (CDI) for support during this study. The authors also thank Buddy Ratner and Jonathan Himmelfarb for thoughtful discussion concerning the role of protein bound uremic toxins in chronic kidney disease and current strategies for targeted PBUT removal.

## References

1. Vanholder, R. C.; De Smet, R. V.;Ringoir, S. M. Assessment of urea and other uremic markers for quantification of dialysis efficacy. Clin. Chem. 38, 1429–1436 (1992).

2. Itoh, Y.; Ezawa, A.; Kikuchi, K.; Tsuruta, Y.; Niwa, T. Protein-bound uremic toxins in hemodialysis patients measured by liquid chromatography/tandem mass spectrometry and their effects on endothelial ROS production. Anal. Bioanal. Chem. 403, 1841–1850 (2012).

3. Vanholder, R.; Schepers, E.; Pletinck, A.; Nagler, E. V.; Glorieux, G. The uremic toxicity of indoxyl sulfate and p-cresyl sulfate: A systematic review. J. Am. Soc. Nephrol. 25, 1897–1907 (2014).

4. Leong, S. C.; Sirich, T. L. Indoxyl sulfate - review of toxicity and therapeutic strategies. Toxins (Basel). 8, (2016).

5. Gryp, T.; Vanholder, R.; Vaneechoutte, M.; Glorieux, G. P-cresyl sulfate. Toxins (Basel). 9, 1–24 (2017).

6. Sirich, T. L.; Aronov, P. A.; Plummer, N. S.; Hostetter, T. H.; Meyer, T. W. Numerous protein-bound solutes are cleared by the kidney with high efficiency. Kidney Int. 84, 585–590 (2013).

7. Sakai, T.; Takadate, A.; Otagiri, M. Characterization of binding site of uremic toxins on human serum albumin. Biol. Pharm. Bull. 18, 1755–1761 (1995).

8. Tao, X.; Thijssen, S.; Kotanko, P.; Ho, C. H.; Henrie, M.; Stroup, E.; Handelman, G. Improved dialytic removal of protein-bound uraemic toxins with use of albumin binding competitors: An in vitro human whole blood study. Sci. Rep. 6, 2–10 (2016).

9. Böhringer, F.; Jankowski, V.; Gajjala, P. R.; Zidek, W.; Jankowski, J. Release of uremic retention solutes from protein binding by hypertonic predilution hemodiafiltration. ASAIO J. 61(1), 55–60 (2015).

10. Krieter, D. H.; Devine, E.; Küorner, T.; Rüth, M.; Wanner, C.; Raine, M.; Jankowski, J.; Lemke, H.D. Haemodiafiltration at increased plasma ionic strength for improved protein-bound toxin removal. Acta Physiol 219, 510–520 (2017).

11. Sandeman, S. R.; Zheng, Y.; Ingavle, G. C.; Howell, C. A.;Mikhalovsky, S. V.; Basnayake, K.; Boyd, O.; Davenport, A.; Beaton, N.; Davies, N. A haemocompatible and scalable nanoporous adsorbent monolith synthesised using a novel lignin binder route to augment the adsorption of poorly removed uraemic toxins in haemodialysis A haemocompatible and scalable nanoporous adsorbent monolith synthesised. Biomed. Mater. 12, 035001 (2017).

12. Sandeman, S. R.; Howell, C. A.; Phillips, G. J.; Zheng, Y.; Boyd, O.; Holt, S.; Mikhalovsky, S. V. An adsorbent monolith device to augment the removal of uraemic toxins during haemodialysis. J. Mater. Sci. Mater. Med. 25, 1589–1597 (2014).

13. Ghuman, J.; Zunszain, P. A.; Petitpas, I.; Bhattacharya, A. A.; Otagiri, M.; Curry, S.; Structural basis of the drug-binding specificity of human serum albumin. J. Mol. Biol. 353, 38–52 (2005).

14. Florens, N.; Yi, D.; Juillard, L.; Soulage, C. O. Using binding competitors of albumin to promote the removal of protein-bound uremic toxins in hemodialysis: Hope or pipe dream? Biochimie 144, 1–8 (2018).

15. Devine, E.; Krieter, D. H.; Rüth, M.; Jankovski, J.; Lemke, H. D. Binding affinity and capacity for the uremic toxin indoxyl sulfate. Toxins (Basel). 6, 416–430 (2014).

16. Case, D. A. Amber 2014. (2014).

17. Humphrey, W.; Dalke, A.; Schulten, K. VMD: Visual molecular dynamics. J. Mol. Graph. 14, 33–38 (1996).

18. Frisch, M. J. et al. Gaussian 09. (2010).

19. Abraham, M. J. et al. Gromacs: High performance molecular simulations through multi-level parallelism from laptops to supercomputers. SoftwareX 1-2, 19–25 (2015).

20. Jorgensen, W. L.; Chandrasekhar, J.; Madura, J. D.; Impey, R. W.; Klein, M. L. Comparison of simple potential functions for simulating liquid water. J Chem Phys 79, 926–935 (1983).

21. Maier, J. A.; Martinez, C.; Kasavajhala, K.; Wickstrom, L.; Hauser, K. E.; Simmerling, C. ff14SB: Improving the accuracy of protein side chain and backbone parameters from ff99SB. J. Chem. Theory Comput. 11, 3696–3713 (2015).

22. Bayly, C. I.; Cieplak, P.; Cornell, W. D.; Kollman, P. A. A well-behaved electrostatic potential based method using charge restraints for deriving atomic charges: The RESP model. J. Phys. Chem. 97, 10269–10280 (1993).

23. Wang, J. M.; Wolf, R. M.; Caldwell, J. W.; Kollman, P. A.; Case, D. A. Development and testing of a general amber force field. J. Comput. Chem. 25, 1157–1174 (2004).

24. Bussi, G.; Donadio, D.; Parrinello, M. Canonical sampling through velocity rescaling. J. Chem. Phys. 126, 014101 (2007).

25. Berendsen, H. J. C.; Postma, J. P. M.; Van Gunsteren, W. F.; Dinola, A.; Haak, J. R. Molecular dynamics with coupling to an external bath. J. Chem. Phys. 81, 3684–3690 (1984).

26. Parrinello, M.; Rahman, A. Polymorphic transitions in single crystals: A new molecular dynamics method. J. Appl. Phys. 52, 7182–7190 (1981).

27. Tribello, G. A.; Bonomi, M.; Branduardi, D.; Camilloni, C.; Bussi, G. PLUMED 2: New feathers for an old bird. Comput. Phys. Commun. 185, 604–613 (2014).

28. Comaniciu, D.; Meer, P. Mean shift: A robust approach toward feature space analysis. IEEE Trans. Pattern Anal. Mach. Intell. 24, 603–619 (2002).

29. Gasparotto, P.; Ceriotti, M. Recognizing molecular patterns by machine learning: An agnostic structural definition of the hydrogen bond. J. Chem. Phys. 141, 1–14 (2014).

30. Gasparotto, P.; Meißner, R. H.; Ceriotti, M. Recognizing local and global structural motifs at the atomic scale. J. Chem. Theory Comput. 14, 486–498 (2018).

31. Rousseeuw, P. J. Silhouettes: a graphical aid to the interpretation and validation of cluster analysis. J. Comput. Appl. Math. 20, 53–65 (1987).

32. Watanabe, H.; Noguchi, T.; Miyamoto, Y.; Kadowaki, D.; Kotani, S.; Nakajima, M.; Miyamura, S.; Ishima, Y.; Otagiri, M.; Maruyama, T. Interaction between two sulfate-conjugated uremic toxins, p-cresyl sulfate and indoxyl sulfate, during binding with human serum albumin. Drug Met. Disp. 40, 1423–1428 (2012).

33. Li, J.; Han, L.; Liu, S.; He, S.; Cao, Y.; Xie, J.; Jia, L. Removal of indoxyl sulfate by water-soluble poly-cyclodextrins in dialysis. Coll. Surf. B Bioint. 164, 406–413 (2018).

34. Limongelli, V.; Bonomi, M.; Parrinello, M. Funnel metadynamics as accurate binding free-energy method. Proc. Natl. Acad. Sci. 110, 6358–6363 (2013).

35. Tiwary, P.; Parinello, M. From metadynamics to dynamics. Phys. Rev. Lett. 111, 230602 (2013).

