## Supplementary material for "Elucidating the molecular interactions between uremic toxins and the Sudlow II binding site of human serum albumin": SI

FIGURES

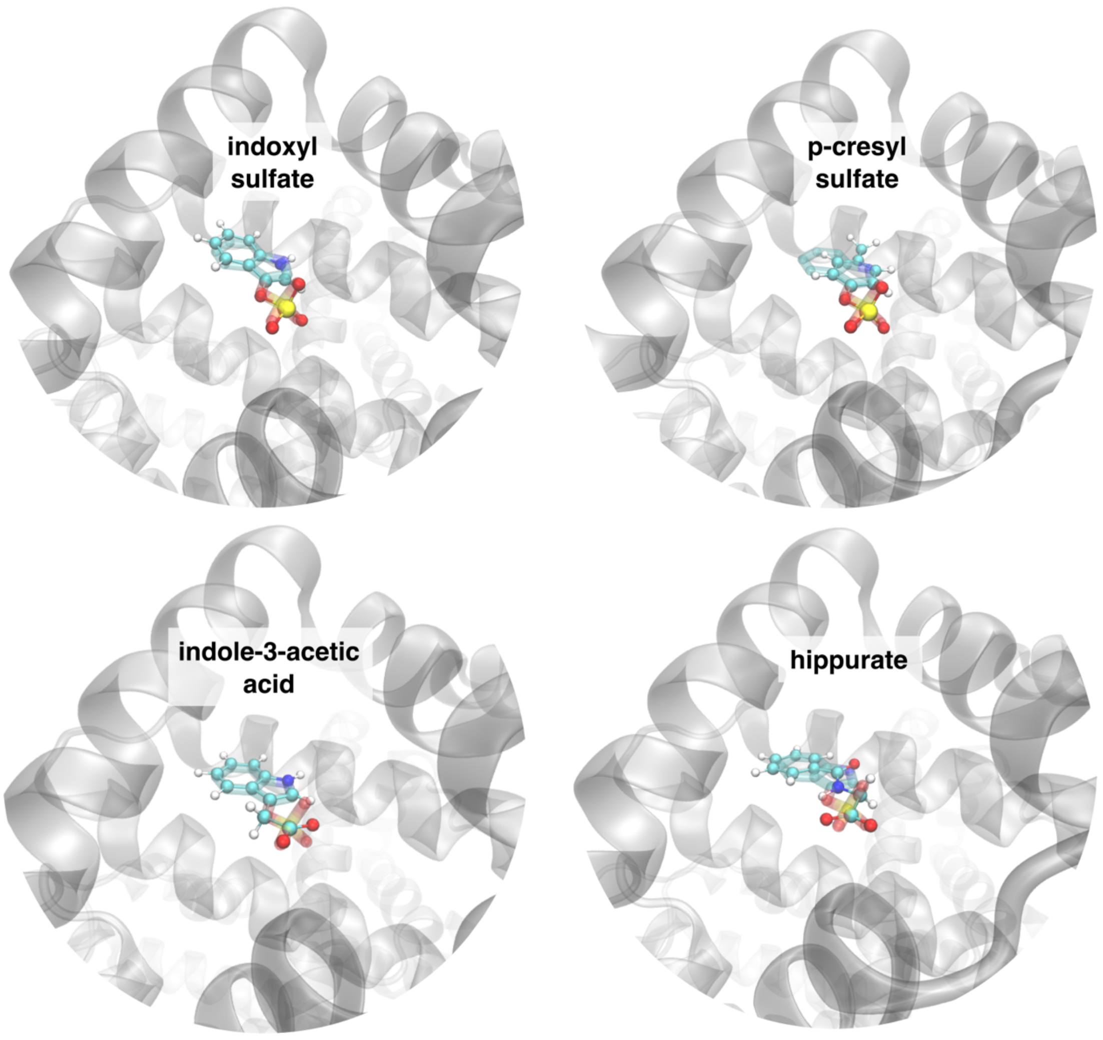


**Figure S1. Binding poses used to initialize (top left) indoxyl sulfate-HSA, (top right) p-cresyl sulfate-HSA, (bottom left) indole-3-acetic acid-HSA, and (bottom right) hippurate-HSA simulations.** The initial protein structure (transparent gray) is taken in all 4 cases from the experimental x-ray structure for IS bound to Sudlow's Site II (PDB ID: 2BXH), and each toxin molecule (opaque ball and stick) was superimposed in place of the experimentally resolved IS molecule (transparent, colored by atom).

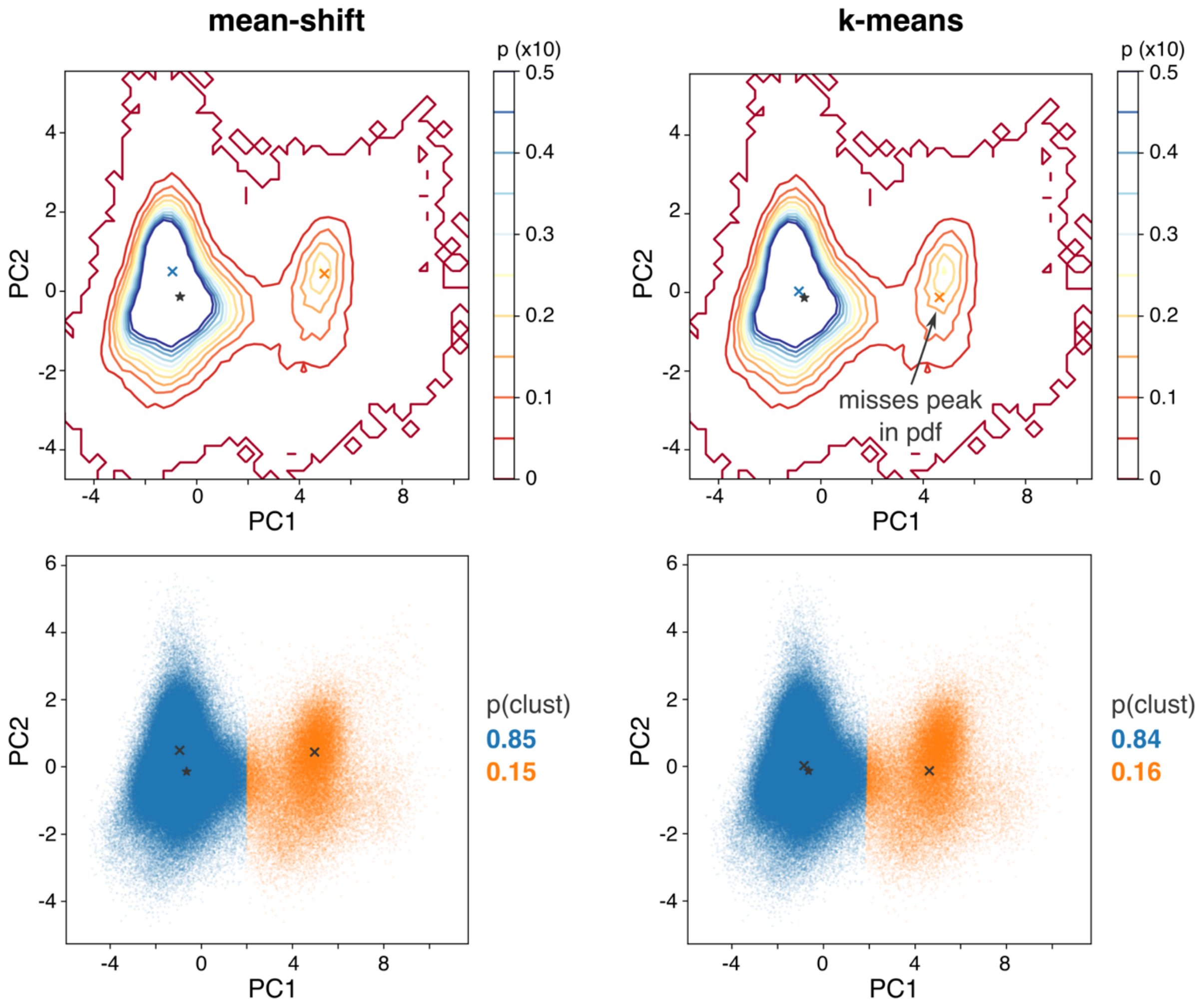


**Figure S2. Comparison of conformation clustering results for IS-HSA complex from the mean-shift and k-means algorithms.** (left) Because mean-shift is a "mode-seeking algorithm", cluster centers converge to peaks on the underlying probability density function. (right) The k-means algorithm tends to converge on clusters with similar spatial extent (here in 2D principal component space) without consideration of the underlying probability density. The centers of the k-means clusters are less representative of metastable binding modes. Ultimately, the dividing line between the two clusters for the IS-HSA complex is insensitive to the clustering method.

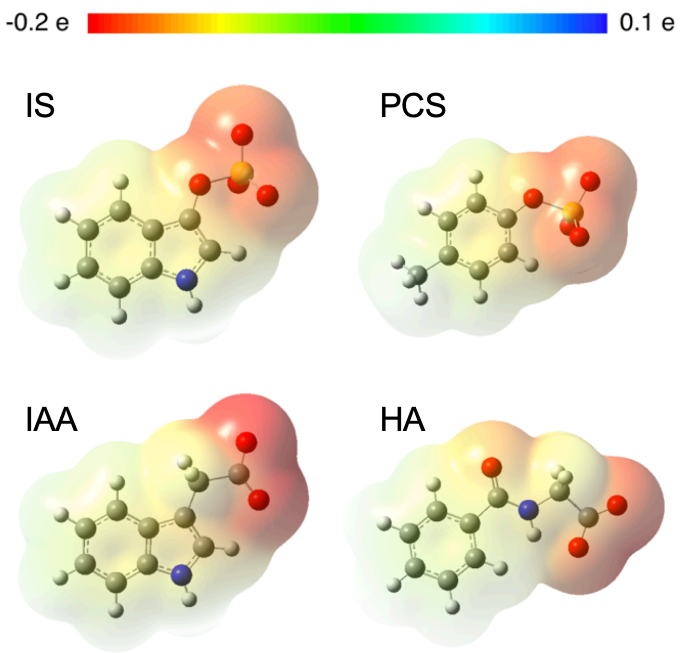


**Figure S3. Electrostatic potential surface visualization for each PBUT, based on point charges from the RESP method.** Darker red represents regions of more negative electrostatic charge, while green and dark blue represent neutral and positive charge, respectively. The electrostatic potential surface here is mapped onto an electron density isosurface at 0.0004 (number density).

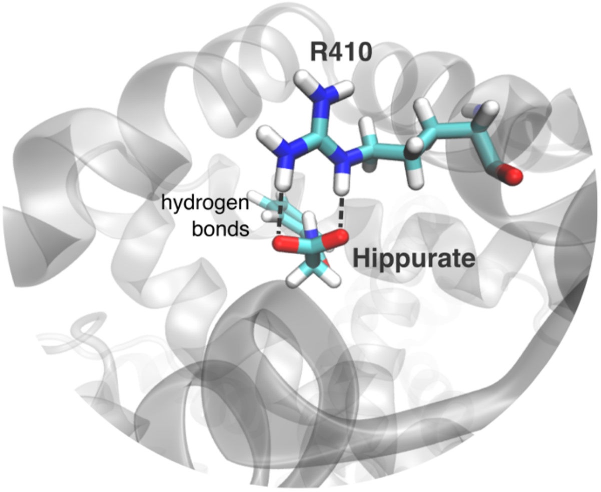


**Figure S4. Representative image of carboxylate-R410 double hydrogen bonded conformation for HA.** This double hydrogen bonded geometry was also observed for IAA-R410 interactions.

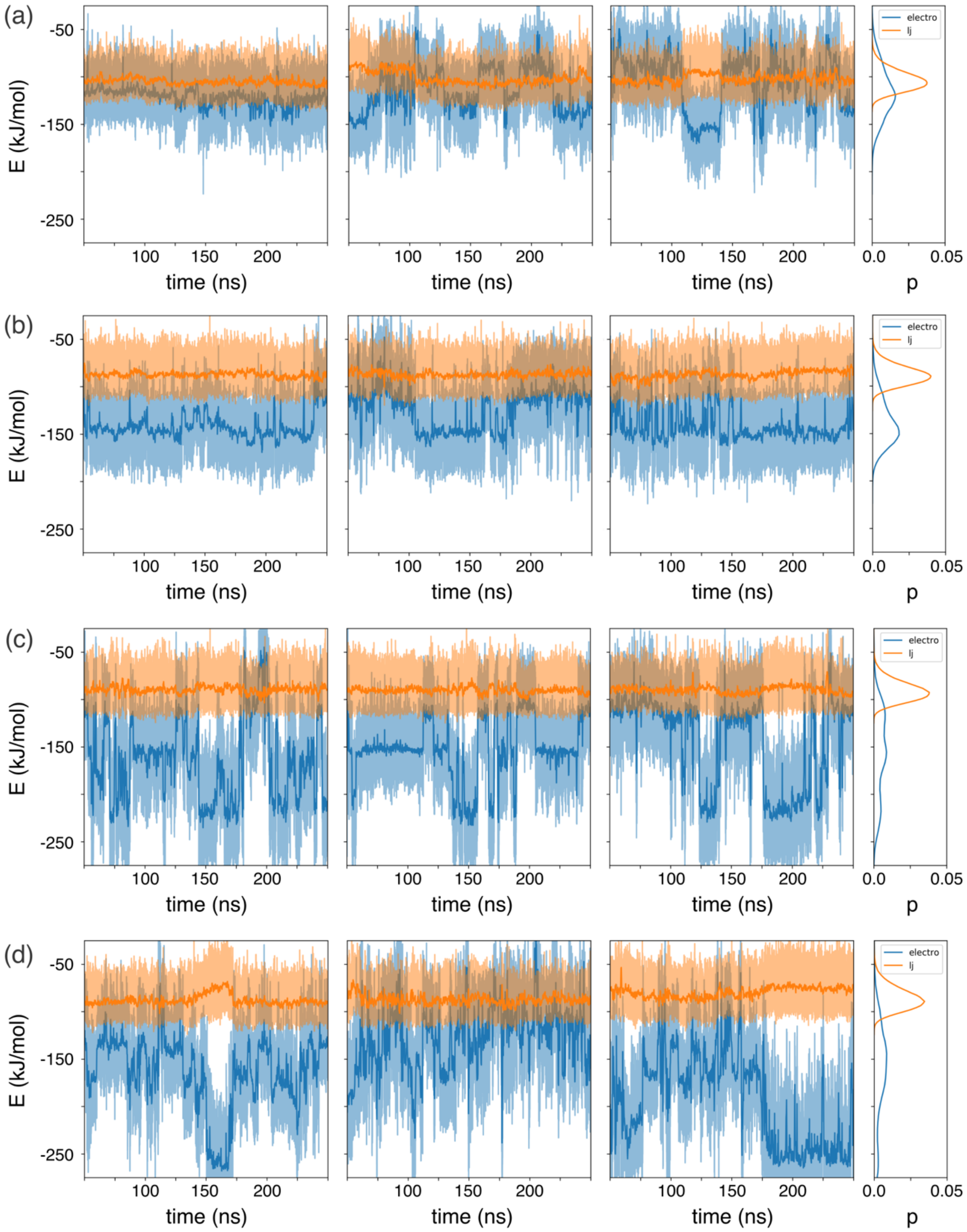


**Figure S5. The protein-toxin interaction energy calculated from MD simulations of HSA in complex with (top) indoxyl sulfate and (bottom) p-cresyl sulfate.** The light blue and light orange shaded areas represent the Coulomb and Lennard-Jones energies, respectively, calculated every 2 ps of simulation time. The dark blue and dark orange lines represent the average Coulomb and Lennard-Jones energies, respectively, over the previous 0.2 ns (i.e. the mean of the last 100 observations at 2 ps intervals). The rightmost panel of each row shows the probability distribution function of each energy for all three simulations of the given toxin based on simple histogramming.

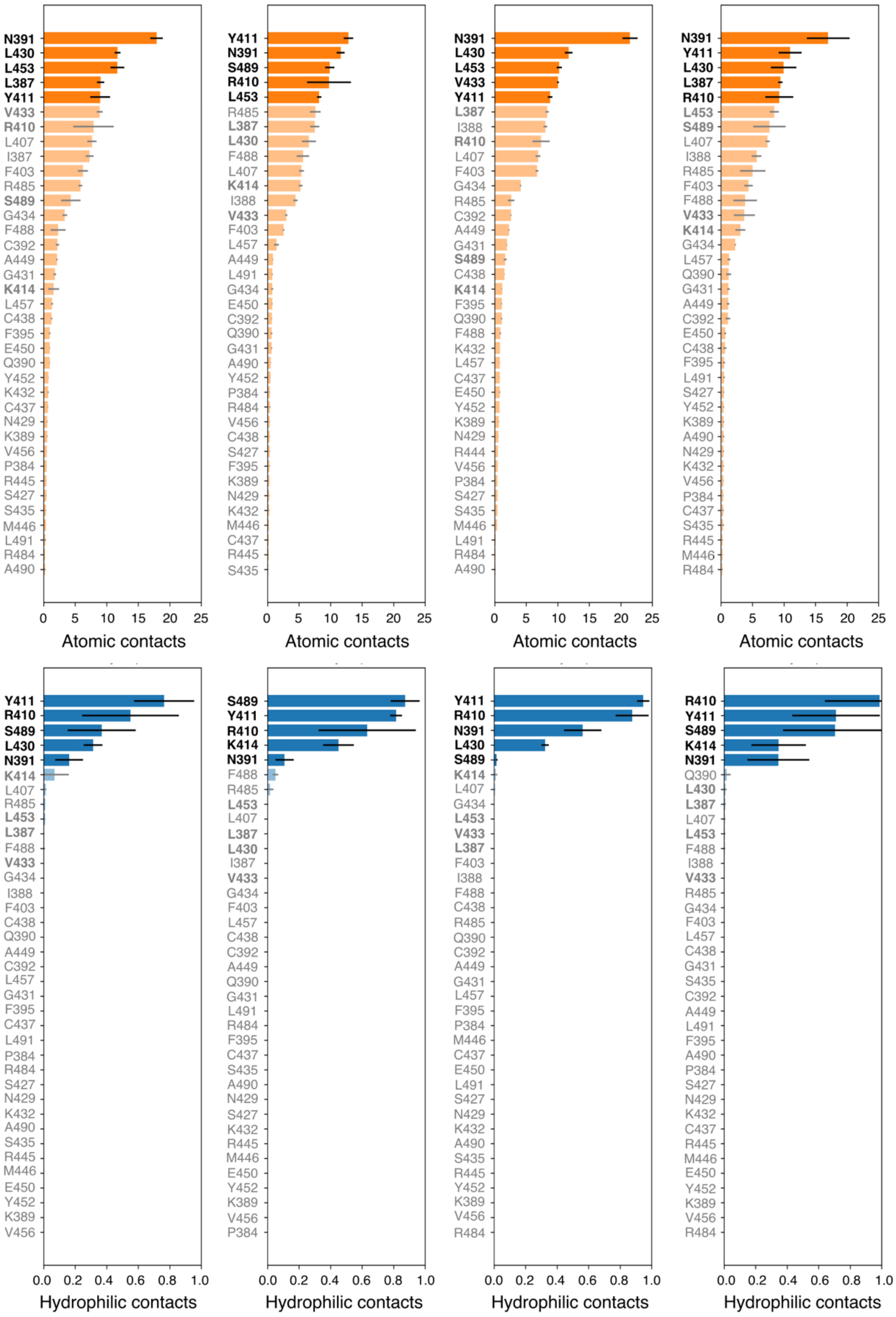


**Figure S6. Average number of atomic and hydrophilic contacts observed in the MD simulations for the 37 residues within 6 Å of IS in the X-ray structure (pdb: 2BXH).** The top 5 residues in each bar plot were added to the set of key residues, which resulted in 9 residues to be considered for further analysis.

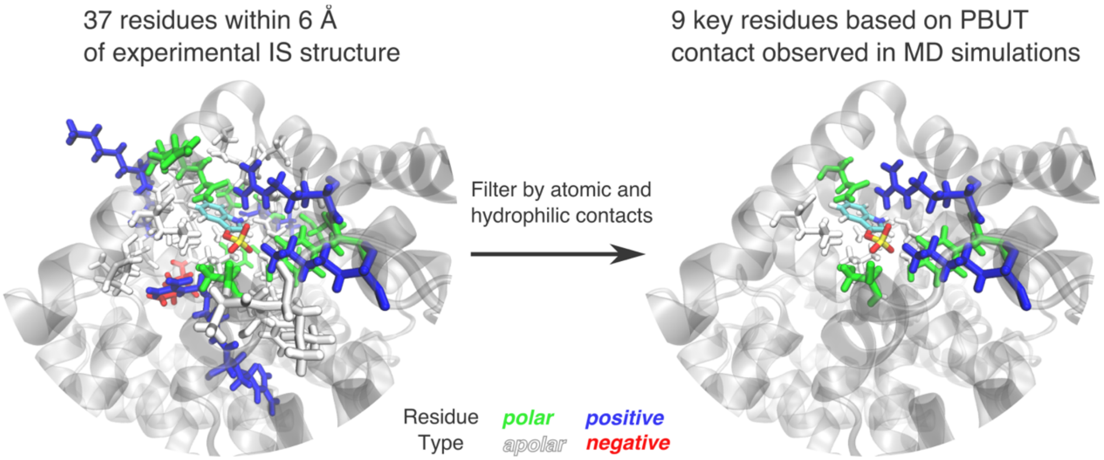


**Figure S7. IS-HSA protein data bank structure (2BXH) with highlighted (left) the 37 residues considered before atomic contact filtering and (right) the 9 residues remaining after filtering by atomic contacts.** Polar, apolar, positive, and negative amino acids are colored green, white, blue, and red, respectively. Indoxyl sulfate is colored by atom type with hydrogen, carbon, oxygen, and sulfur atoms colored white, cyan, red, and yellow, respectively.

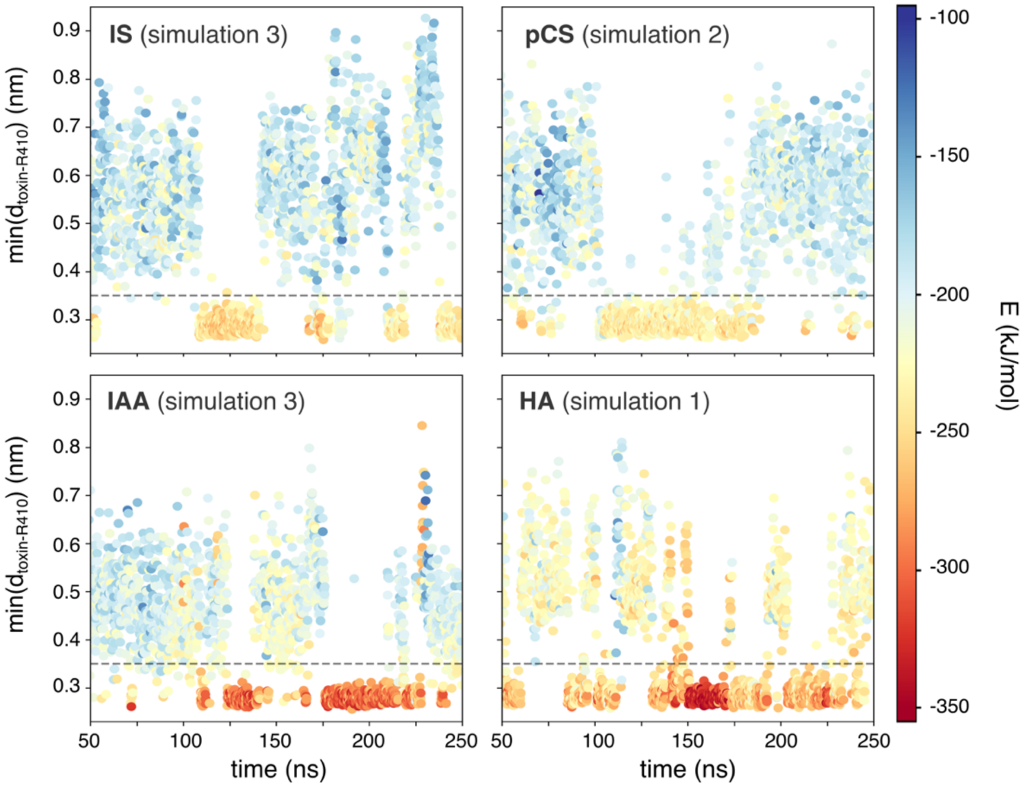


**Figure S8. The minimum distance between each PBUT and R410 heavy atoms vs. simulation time, and it's relation to the overall protein-toxin interaction energy.** Each point is colored based on the overall protein-toxin interaction energy, ranging from dark red (lower energy) to dark blue (higher energy). The dashed horizontal line at 0.35 nm represents a putative cutoff for toxin-R410 hydrogen bonds. The simulation that best captures the transient nature of the R410 complex for each PBUT is pictured, although the breaking/forming of this salt bridge was observed at least once in each production simulation.

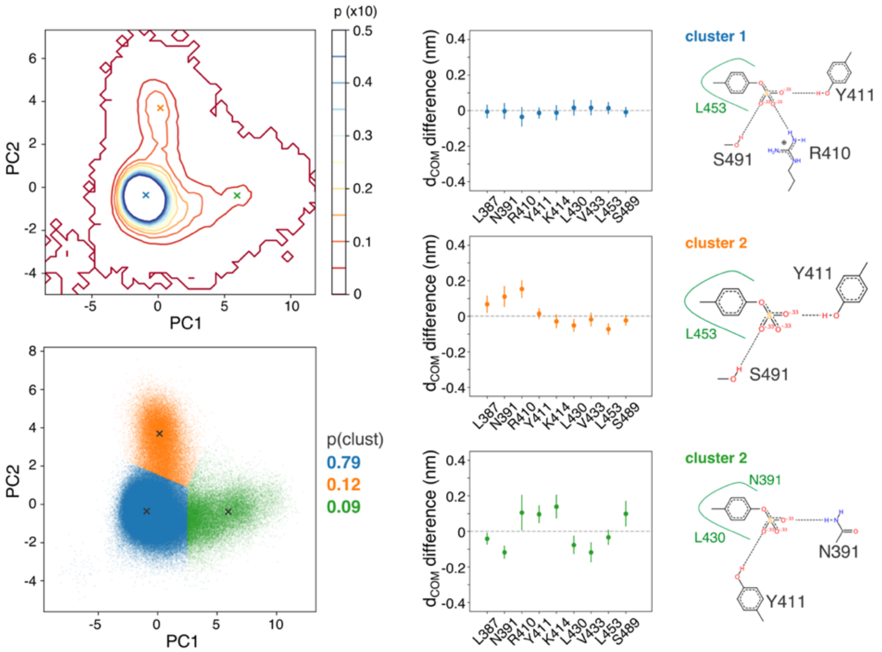


**Figure S9. PCA and clustering results for the pCS-HSA complex. (top left)** Contour plot of the 2D pdf for the pCS-HSA complex, created from a 2D histogram of the PC values for each frame of the MD simulations, with a probability step of 0.005 between contour lines. Blue, orange, and green 'x' symbols mark the center of each mode identified with mean-shift. **(bottom left)** Scatter plot of the first 2 PC values for each MD frame, colored by the cluster assigned by mean-shift. **(center column)** The difference between the average pCS-residue center of mass distance for all points in each cluster and the overall average pCS-residue center of mass distance (all points in all clusters). **(right column)** A PoseView representation of the central structure of each cluster (the MD frame with PC values nearest the 'x' symbols).

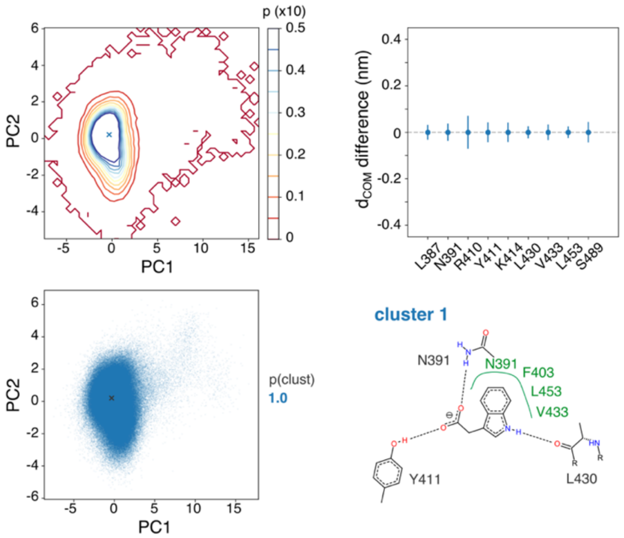


**Figure S10. PCA and clustering results for the IAA-HSA complex. (top left)** Contour plot of the 2D pdf for the IAA-HSA complex, created from a 2D histogram of the PC values for each frame of the MD simulations, with a probability step of 0.005 between contour lines. Blue 'x' symbol marks the center of the lone binding mode identified with mean-shift. **(bottom left)** Scatter plot of the first 2 PC values for each MD frame. **(top right)** The difference between the average IAA-residue center of mass distance for all points in the cluster and the overall average IAA-residue center of mass distance (all points in all clusters). **(bottom right)** A PoseView representation of the central structure of the lone cluster (the MD frame with PC values nearest the 'x' symbol).

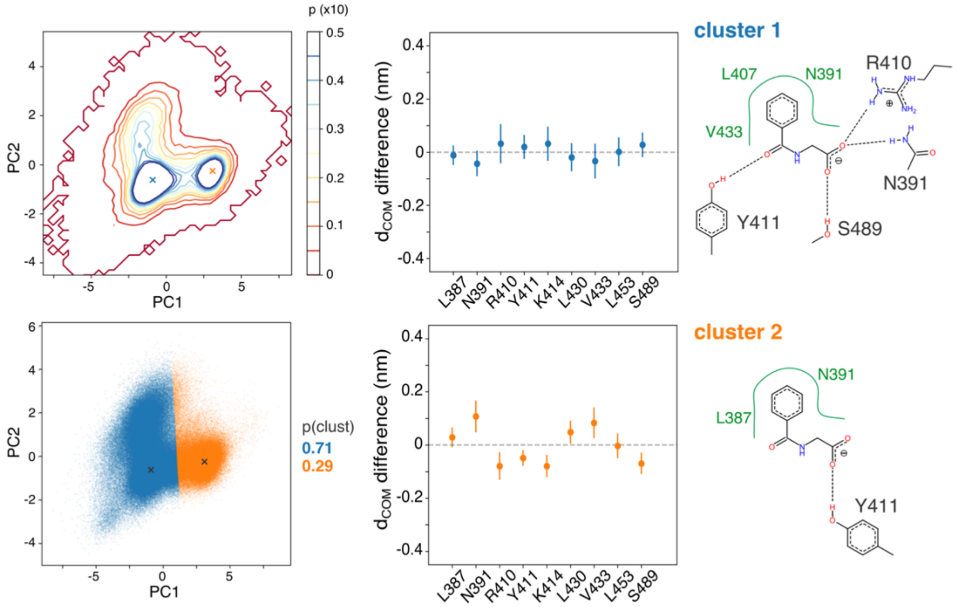


**Figure S11. PCA and clustering results for the hippurate-HSA complex. (top left)** Contour plot of the 2D pdf for the HA-HSA complex, created from a 2D histogram of the PC values for each frame of the MD simulations, with a probability step of 0.005 between contour lines. Blue and orange 'x' symbols mark the center of each mode identified with mean-shift. **(bottom left)** Scatter plot of the first 2 PC values for each MD frame, colored by the cluster assigned by mean-shift. **(center column)** The difference between the average HA-residue center of mass distance for all points in each cluster and the overall average HA-residue center of mass distance (all points in all clusters). **(right column)** A PoseView representation of the central structure of each cluster (the MD frame with PC values nearest the 'x' symbols in **(a-b)**.

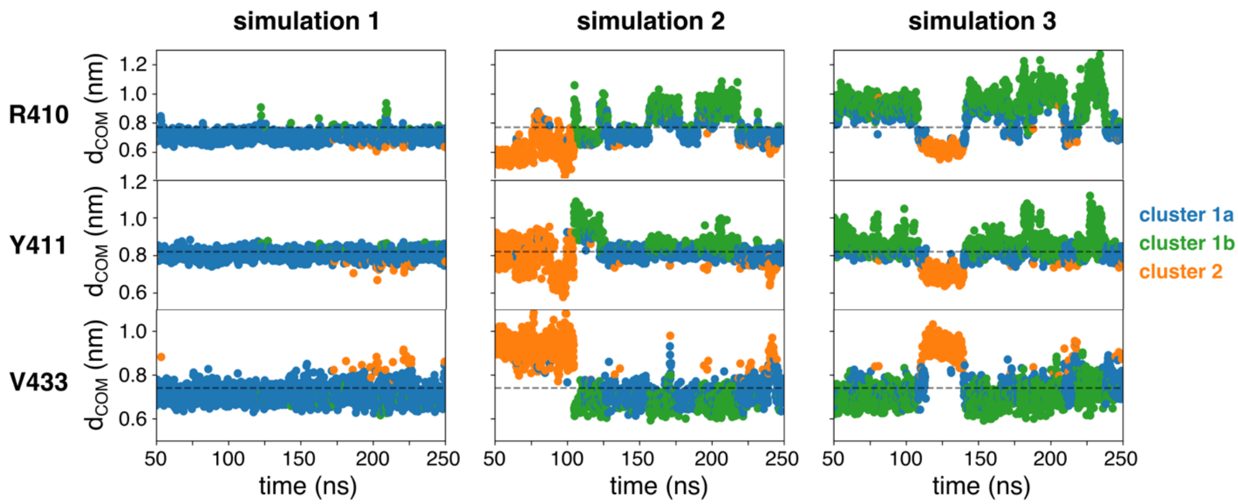


**Figure S12. Three residue-IS distances vs. simulation time, and their relation to the structural cluster id assigned by k-means clustering (k = 3).** For each of the three MD simulations of the IS-HSA complex, the (top) IS-R406, (middle) IS-Y407, and (bottom) IS-V429 center of mass distances are plotted at 0.1 ns intervals between 50 and 250 ns. Colors are assigned according to the cluster id assigned by k-means, named (blue) cluster 1a, (green) cluster 1b, and (orange) cluster 2. The horizontal dashed line in each row of plots represents the overall mean of the given residue-toxin distance.

TABLES

**Table S1. Coefficients of the linear transformation to the first 2 principal components calculated for the IS-HSA and pCS-HSA complex.**

| | indoxyl sulfate | | | p-cresyl sulfate | | | indole-3-acetic acid | | | hippurate | | |
|---|---|---|---|---|---|---|---|---|---|---|---|---|
| | **PC1 (0.57)** | **PC2 (0.15)** | PC3 (0.09) | **PC1 (0.46)** | **PC2 (0.24)** | PC3 (0.09) | **PC1 (0.28)** | **PC2 (0.17)** | PC3 (0.12) | **PC1 (0.48)** | **PC2 (0.15)** | PC3 (0.12) |
| L383 | 0.03 | 0.67 | 0.69 | - 0.16 | 0.51 | 0.34 | 0.13 | - 0.55 | - 0.01 | 0.25 | - 0.01 | - 0.60 |
| N387 | 0.37 | 0.20 | 0.11 | - 0.29 | 0.47 | 0.05 | 0.41 | - 0.37 | - 0.06 | 0.41 | - 0.10 | - 0.25 |
| R406 | - 0.28 | - 0.40 | 0.51 | 0.28 | 0.39 | - 0.47 | - 0.36 | 0.26 | 0.42 | - 0.36 | - 0.22 | 0.13 |
| Y407 | - 0.30 | - 0.32 | 0.35 | 0.42 | 0.03 | - 0.05 | - 0.31 | - 0.24 | 0.60 | - 0.38 | - 0.17 | - 0.12 |
| K410 | - 0.40 | 0.11 | -0.09 | 0.42 | - 0.17 | 0.20 | - 0.46 | - 0.16 | 0.12 | - 0.35 | - 0.37 | 0.19 |
| L426 | 0.38 | - 0.02 | -0.04 | - 0.31 | - 0.28 | 0.44 | 0.16 | 0.39 | 0.09 | 0.30 | - 0.41 | 0.45 |
| V429 | 0.41 | - 0.05 | 0.05 | - 0.43 | - 0.06 | - 0.04 | 0.44 | 0.23 | 0.25 | 0.37 | - 0.04 | 0.41 |
| L449 | 0.33 | - 0.32 | 0.18 | - 0.21 | - 0.48 | - 0.43 | - 0.14 | 0.42 | - 0.26 | - 0.01 | 0.74 | 0.36 |
| S485 | - 0.33 | 0.37 | -0.27 | 0.36 | - 0.13 | 0.48 | - 0.38 | - 0.16 | - 0.56 | - 0.38 | 0.23 | - 0.11 |

***first 2 PCs used for pdf visualizations and clustering**

**Table S2. Silhouette scores for mean-shift clustering with various bandwidths.**

| | **IS** | **pCS** | IAA | HA |
|---|---|---|---|---|
| 0.5 | 0.20 | 0.30 | 0.18 | 0.30 |
| 0.6 | 0.32 | 0.33 | 0.24 | 0.44 |
| 0.7 | 0.57 | 0.40 | 0.37 | 0.46 |
| 0.8 | 0.58 | 0.50 | 0.32 | 0.46 |
| 0.9 | 0.61 | **0.55** | 0.37 | 0.46 |
| 1.0 | 0.60 | 0.55 | 0.45 | 0.46 |
| 1.1 | **0.64** | 0.55 | 0.60 | **0.52** |
| 1.2 | 0.64 | 0.55 | 0.54 | 0.52 |
| 1.3 | 0.64 | 0.50 | 0.64 | 0.52 |
| 1.4 | 0.64 | 0.50 | 0.67 | 0.52 |
| 1.5 | 0.64 | 0.50 | 0.56 | 0.52 |
| 1.6 | 0.64 | 0.50 | 0.67 | 0.52 |
| 1.7 | 0.64 | 0.50 | 0.66 | 0.52 |
| 1.8 | 0.64 | 0.50 | 0.67 | 0.52 |
| 1.9 | 0.64 | 0.50 | **0.69** | 0.52 |
| 2.0 | 0.64 | - | 0.66 | 0.52 |

***bold values selected for clustering**

**Table S3. Average number of hydrogen bonds for each complex.**

| Simulation ID | IS | pCS | IAA | HA |
|---|---|---|---|---|
| 1 | 2.84 | 3.21 | 3.19 | 3.71 |
| 2 | 2.40 | 2.64 | 3.13 | 2.97 |
| 3 | 2.53 | 2.99 | 3.25 | 4.00 |
| mean | 2.59 (0.184) | 2.95 (0.238) | 3.19 (0.050) | 3.56 (0.436) |
